# The inner nuclear membrane-associated protein Csa1 regulates lipid droplet biogenesis and nuclear shape in a Sir2-dependent manner

**DOI:** 10.1101/2025.07.25.666785

**Authors:** Lauren Carson, Hui Jin, Hong-Guo Yu

**Affiliations:** Department of Biological Science, Florida State University, Tallahassee FL 32306

**Author notes:** Correspondence to: Hong-Guo Yu.

## Abstract

The nuclear envelope, a double-membrane structure consisting of the inner nuclear membrane (INM) and the outer nuclear membrane, encloses the nucleus. In response to cellular cues, lipid droplets are formed at the INM, which plays a key role in membrane homeostasis and energy storage. However, the signaling pathway that regulates the formation of nuclear lipid droplets remains to be elucidated. Here we show that in budding yeast, the INM-localized protein Csa1 regulates the nuclear localization of the acyltransferase Lro1, which converts diacylglycerol to triacylglycerol, leading to lipid droplet formation. Csa1 harbors a bipartite nuclear localization signal for nuclear sorting, along with an amphipathic helix that anchors it to the INM. Upon nutrient depletion and at the onset of meiosis, both Csa1 and Lro1 are enriched at a distinct INM subdomain that abuts the nucleolus, which we term the n-INM. We show that Csa1 is essential for targeting Lro1 to the n-INM region. In the absence of Csa1, Lro1 is dispersed along the nuclear periphery. The deacetylase activity of Sir2 is required for the n-INM enrichment of both Csa1 and Lro1. In the absence of either Csa1 or Sir2, lipid droplet formation is compromised. Our findings uncover an unexpected Sir2-dependent Csa1-Lro1 signaling pathway at the INM that regulates lipid droplet biogenesis and nuclear envelope remodeling.

## Introduction

The nuclear envelope consists of two membranes, the inner nuclear membrane (INM) and the outer nuclear membrane (ONM). The ONM is continuous with the endoplasmic reticulum (ER), while the INM forms a specialized cellular compartment that interacts with the nucleoplasm and the genome. The INM and ONM are joined at the nuclear pore complexes, openings on the nuclear envelope that regulate transport between the nucleoplasm and the cytoplasm (Ungricht and Kutay, 2017; Mannino and Lusk, 2022). Recent research has uncovered that the INM, unexpectedly, recruits numerous ER-associated enzymes involved in lipid metabolism and is actively engaged in the synthesis of lipid droplets (Witkin et al., 2012; Barbosa et al., 2015; Romanauska and Kohler, 2018; Barbosa et al., 2019; Romanauska and Kohler, 2021; Soltysik et al., 2021; Foo et al., 2023). The INM is thought to maintain high levels of diacylglycerol (DAG), a key precursor for triacylglycerol (TAG), which is the primary neutral lipid found in lipid droplets (Romanauska and Kohler, 2018; Foo et al., 2023; Lee et al., 2023). While both cellular signaling and gene network changes are necessary for the synthesis of nuclear lipid droplets (Romanauska and Kohler, 2021; Fujimoto, 2024; Romanauska et al., 2024), the signaling pathways that control TAG production at the nuclear envelope remain to be further elucidated. Additionally, the biological function of nuclear lipid droplet formation is largely unexplored.

The ER-associated acyltransferase Lro1 in budding yeast catalyzes the conversion of DAG to TAG, facilitating the formation of lipid droplets that serve as cytoplasmic hubs for energy storage and membrane remodeling (Oelkers et al., 2000; Lysyganicz et al., 2025). Lro1 is a type II single transmembrane domain protein, with its enzymatic C-terminus located in the ER lumen (Choudhary et al., 2011; Barbosa et al., 2019). In budding yeast, lipid droplet formation is mediated by four acyltransferases — Lro1, Dga1, Are1, and Are2— which are functionally redundant (Oelkers et al., 2002; Sandager et al., 2002). Together, these four acyltransferases are critical for maintaining lipid homeostasis by regulating the levels of phosphatidic acid, the precursor to membrane lipids, and DAG (Ruggles et al., 2013). Notably, Lro1 is the only acyltransferase known to be imported into the nucleus (Barbosa et al., 2015; Barbosa et al., 2019). In starved yeast cells, Lro1 accumulates at a distinct INM subdomain that abuts the nucleolus (Barbosa et al., 2019), a membraneless subnuclear compartment formed by the repeated rDNA gene array and its associated proteins and RNAs. This specific INM region is referred to as the nucleolus-associated INM (n-INM), as detailed below.

The n-INM region in budding yeast is known to contain a lower density of nuclear pore complexes and lacks the basket nucleoporins (Strambio-de-Castillia et al., 1999; King et al., 2023). During mitotic arrest, the nuclear envelope expands specifically at the n-INM region, forming what is referred to as a nuclear flare (Witkin et al., 2012; Walters et al., 2014). Notably, overexpression of Lro1 suppresses the formation of the nuclear flare (Barbosa et al., 2019), demonstrating that both the localization of Lro1 to the n-INM and its enzymatic activity are necessary for the synthesis of nuclear lipid droplets. While the cell cycle-dependent nature of nuclear flare formation has been well established, the mechanisms that restrict this expansion to the n-INM region of the nuclear envelope are not fully understood.

In response to nutrient depletion, the repeated rDNA gene array in budding yeast becomes highly compacted, and rRNA transcription is significantly reduced (Smith et al., 2009). Silencing of rDNA gene expression depends on the enzymatic activity of Sir2, a conserved NAD+-dependent histone deacetylase (Imai et al., 2000; Landry et al., 2000; Smith et al., 2000). The recruitment of Sir2 to the rDNA/nucleolus requires the RENT complex, which consists of Cdc14, Net1, and Sir2, and regulates rDNA silencing and mitotic exit (Straight et al., 1999; Huang and Moazed, 2003). A wealth of research has demonstrated that Sir2 controls key rDNA processes, including replication, transcription, and recombination, making it a critical regulator of cellular homeostasis and aging (Gottlieb and Esposito, 1989; Kaeberlein et al., 1999; Lin et al., 2000). However, it remains unknown whether nucleolus-associated Sir2 is involved in the regulation of nuclear lipid droplet formation.

We show here that Csa1 accumulates at the n-INM in a Sir2-dependent manner in response to nutrient depletion and at the onset of meiosis, the latter of which requires both nitrogen starvation and glucose depletion in budding yeast. Csa1 is a paralog of the yeast spindle pole body component Nbp1 and localizes to the yeast nuclear periphery (Kupke et al., 2011; Ord et al., 2020). Like Nbp1, Csa1 is also present at the spindle pole body during anaphase (see below) and is thought to be a substrate of the polo-like kinase Cdc5 (Ord et al., 2020). However, the role of Csa1 in nuclear envelope dynamics remains unknown.

Using yeast genetics and fluorescence microscopy, we have determined that Csa1 contains a bipartite nuclear localization signal and is an INM-associated membrane protein. Importantly, Csa1 is essential for restricting Lro1 to the n-INM region. Additionally, we show that the deacetylase activity of Sir2 is required for the accumulation of both Csa1 and Lro1 at the n-INM. Our findings uncover an unexpected Sir2-dependnent Csa1-Lro1 signaling pathway at the nuclear periphery that regulates both lipid droplet biogenesis and nuclear envelope remodeling.

## Results

### Csa1 localizes to the INM and concentrates at the n-INM

We seek to understand the mechanisms of nuclear envelope remodeling in budding yeast. A small-scale cytological screen of nuclear envelope-associated proteins tagged with C-terminal GFP revealed that Csa1 accumulated in a subdomain of the nuclear periphery during early meiosis and in nutrient-depleted yeast cells (Fig. 1). To determine whether Csa1 localized to the inner, outer, or both nuclear membranes, we used the bimolecular fluorescence complementation assay (Fig. 1A). This technique involves splitting GFP into two fragments: N-terminal GFP_1-10_ and C-terminal GFP_11_ (Smoyer et al., 2016). When these two sections of GFP are present in the same cellular compartment, they can reconstitute a functional fluorescent molecule (Fig. 1A). Expression of Csa1 fused to GFP1-10 (Csa1-GFP1-10) in the presence of the nucleoplasm reporter (GFP11-mCherry-Pus1) and the nuclear envelope/ER reporter (GFP11-mCherry-Scs2TM) consistently demonstrated Csa1 localization to the nuclear periphery, but not to the cortical ER. (Fig. 1B). Conversely, Csa1-GFP_1-10_ failed to interact with the cytoplasm reporter (GFP_11_-mCherry-Hxk1) or the endoplasmic reticulum lumen reporter (Fig. 1B). Based on these findings, we conclude that Csa1 is an inner nuclear membrane-associated protein.

**Figure 1.**
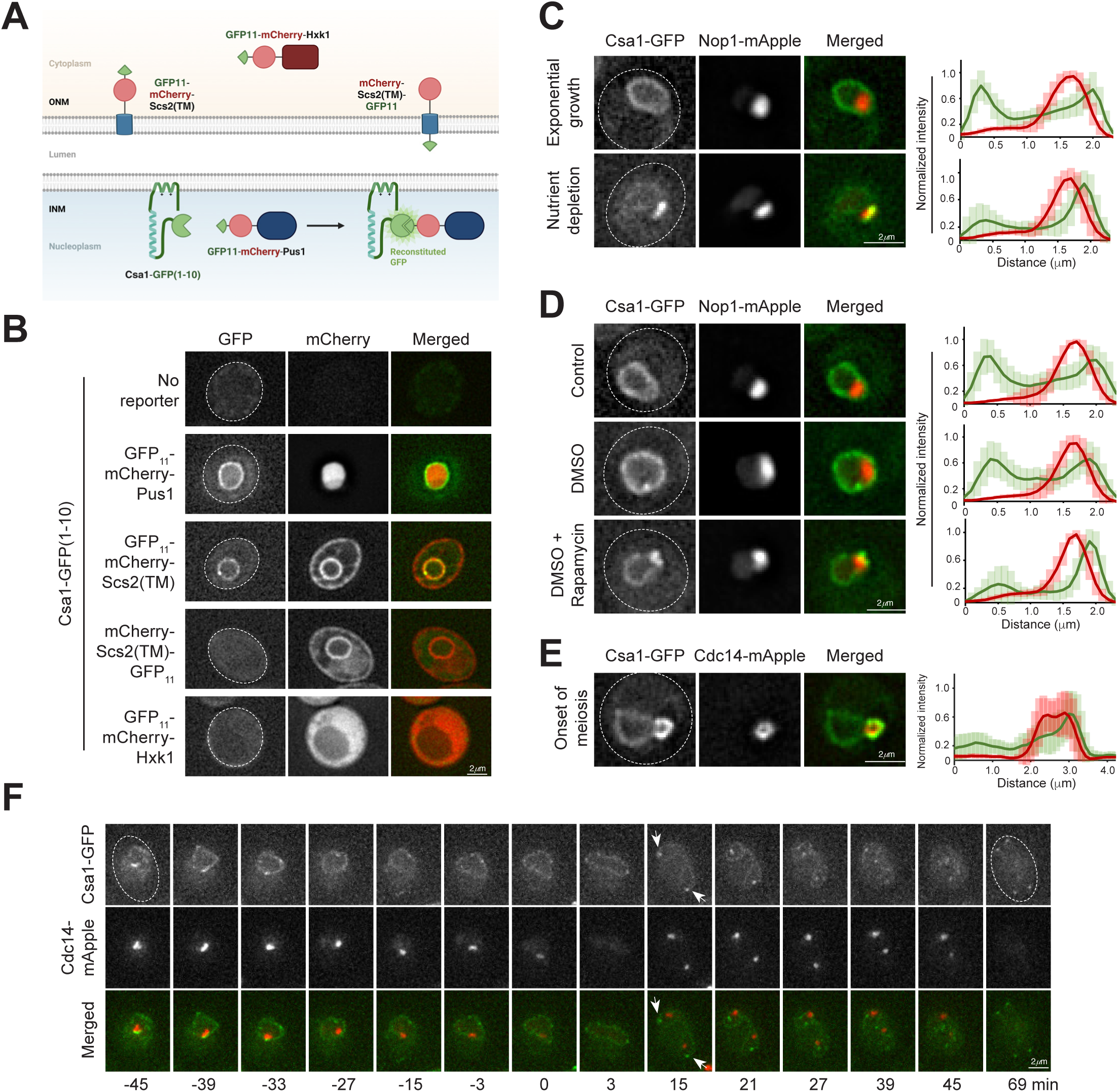
Csa1 localizes to the INM and concentrates at the n-INM region in response to starvation, rapamycin treatment, and the onset of meiosis. (**A**) Schematics of the bimolecular fluorescence complementation assay using split GFP fragments to show Csa1’s association with the inner nuclear membrane (INM). ONM, outer nuclear membrane. (**B**) Representative images show yeast cells expressing Csa1-GFP(1-10) and GFP11 fused to four different reporters: GFP11-mCherry-Pus1 (nucleoplasm), GFP11-mCherry-Scs2(TM) (nuclear envelope and endoplasmic reticulum (ER) membrane), GFP11-Scs2(TM)-mCherry (lumen of the nuclear envelope and ER), and GFP11-mCherry-Hxk1 (cytoplasm). TM, transmembrane domain. Dotted ovals outline cell shape. (**C**) Representative images show Csa1-GFP localization in exponentially growing and nutrient-depleted yeast cells. Nop1-mApple serves as a marker for the nucleolus. Fluorescence intensities along a 2.5 μm line across the nucleus are shown to the right (n=25). Red, Nop1-mApple; green, Csa1-GFP. Error bars represent standard deviation. (**D**) Representative images show Csa1-GFP localization in cells treated with rapamycin for 2 hours. DMSO serves as a vehicle control. Line scans of fluorescence intensity are shown to the right as in **C** (n=25). (**E**) Csa1-GFP localization in a cell staged at the onset of meiosis using *P_DDI2_-IME1*. Cdc14-mApple marks the nucleolus. Note that the nucleolus protrudes outward, and Cdc14 forms a torus shape. Fluorescence intensities along a 5 μm line across the cell are shown to the right (n=25). At least three biological replicates were performed in the experiments presented in **B**-**E**. (**F**) Time-lapse fluorescence microscopy showing Csa1-GFP localization during yeast meiosis. Time zero refers to the onset of anaphase I, when Cdc14 is released from the nucleolus. Arrows indicate the accumulation of Csa1 at the yeast spindle pole body. Scale bars, 2μm.

Next, we investigated the physiological conditions that govern the localization of Csa1. At the beginning of meiosis, Csa1-GFP accumulated in a nuclear subdomain (Fig. 1, see below), which is triggered by nutrient depletion. To understand Csa1 localization during vegetative growth and in starved conditions, including nitrogen-deficient media (SD-N), and during a post-diauxic shift (PDS), we examined Csa1 localization in exponentially growing cells, in SD-N media, and in PDS cells. In exponentially growing cells, Csa1 localized relatively evenly to the nuclear periphery. However, when yeast cells were shifted to SD-N media or in PDS cells, Csa1 was observed to concentrate to the INM that abutted the nucleolus, which was marked by Nop1. (Fig. 1C). We therefore refer to this nucleolus-associated INM as n-INM. Consistent with the idea that the concentration of Csa1 to the n-INM region depends on nutrient availability, treatment with the TORC1 pathway inhibitor rapamycin also resulted in the concentration of Csa1 to the n-INM (Fig. 1D). Finally, we confirmed that Csa1 was concentrated to the n-INM at the beginning of meiosis (Figs. 1E and 1F). The transcription factor Ime1 activates early meiotic genes (Kassir et al., 1988), we therefore constructed a cyanamide-inducible *IME1* allele (*P_DDI2_-IME1*) to arrest yeast cells at early meiosis. Addition of cyanamide allowed yeast cells to resume and complete meiosis without any noticeable defects (our unpublished data). In early meiosis, with or without Ime1 block, the nucleolus formed a sphere-like structure, protruding outward from the rest of the nucleus (Fig. 1E and see below Fig. 4). We found that Csa1 was highly enriched at the interface between this protrusion and the remainder of the nucleus (Fig. 1E and see below Fig. 4). Taken together, our findings identify Csa1 as an inner nuclear membrane protein that concentrates to the n-INM in response to starvation, TORC1 inactivation, and at the beginning of meiosis.

### Csa1 contains a bipartite nuclear localization signal at its N-terminus

We hypothesized that Csa1 possesses a nuclear localization signal for nuclear import. To identify this sorting signal, we created a series of point and deletion mutations in Csa1 based on structural predictions from AlphaFold and sequence similarities to Nbp1 (Fig. 2A and supplemental Fig. S1). The N-terminus of Csa1, including amino acids 1 to 30, was sufficient for nuclear targeting, as evidenced by the concentration of this fragment within the yeast nucleus (Figs. 2B and 2C). Conversely, mutating a cluster of basic residues (Arg11, Lys12, Arg14, Lys15) to Ala in Csa1-(1-30)-NLS1-4A-GFP resulted in cytoplasmic localization, indicating the presence of a nuclear localization sequence (NLS) within this region, which we termed NLS1 (Fig. 2A and 2C). However, this mutation did not completely abolish the nuclear localization of the full-length Csa1 protein (Fig. 2D). Consistent with this observation, the fragment 31-221 of Csa1 was sorted to the nucleus and predominantly localizes to the nuclear periphery (Fig. 2B). These findings suggest that Csa1 possesses multiple nuclear localization sequences.

**Figure 2.**
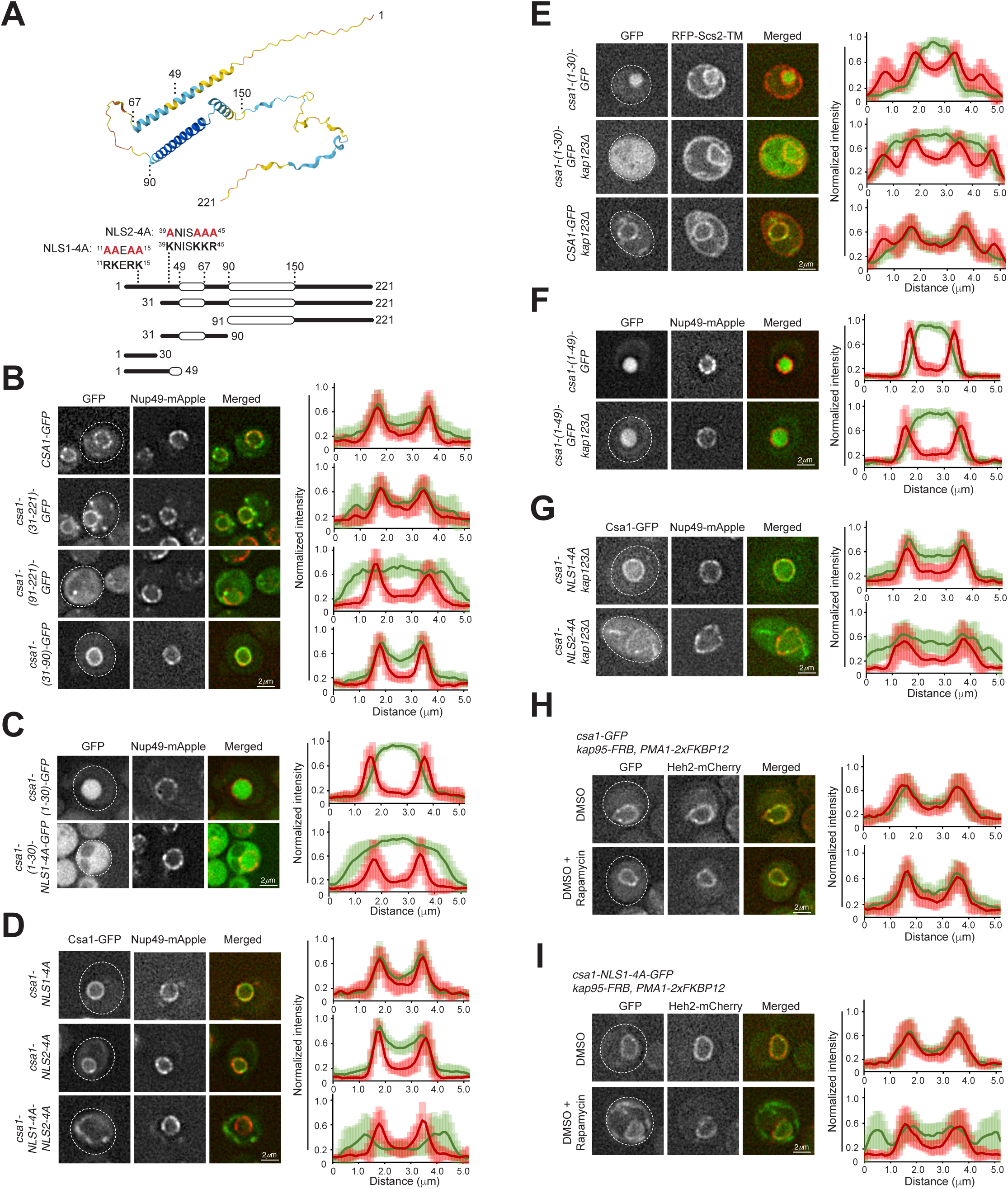
Csa1 contains a bipartite nuclear localization signal at its N-terminus. (**A**) Top: Domain organization of Csa1 predicted by AlphaFold. Bottom: schematics of selected deletion constructs used in this study. Numbers indicate amino acid positions. Amino acids 11 to 15 contain predicted nuclear localization sequence 1 (NLS1), while 39 to 45 contain proposed NLS2. Point mutations of NLS1-4A and NLS2-4A are shown in red. (**B**) The Csa1 fragment from amino acids 1 to 90 is essential for nuclear localization. The expression of the N-terminal deletion constructs was under the control of the *DDI2* promoter. Nup49-mApple marks the nuclear periphery. Fluorescence intensities along a 5 μm line across the cell are shown to the right (n=25). Error bars represent standard deviation. (**C**) The Csa1 fragment from amino acids 1 to 30 contains NLS1. The expression of *csa1-(1-30*) and *csa1-(1-30)-NLS1-4A* was under the control of the endogenous *CSA1* promoter. Note that Csa1-(1-30)-GFP is concentrated in the nucleoplasm, while the NLS1-4A mutation abolishes nuclear confinement of Csa1-(1-30). Line scans of fluorescence intensity are shown on the right as in **B** (n=25). (**D**) Double mutation of NLS1 and NLS2 abolishes Csa1’s concentration at the nuclear envelope. The expression of the mutants was under the control of the endogenous *CSA1* promoter. Line scans of fluorescence intensity are shown on the right as in **B** (n=25). (**E**) Kap123 is required for nuclear import of Csa1-(1-30). Expression of *csa1-(1-30)-GFP* and *CSA1-GFP* was under the control of the endogenous *CSA1* promoter. RFP-Scs2-TM marks the peripheral and cortical ER membranes. Line scans of fluorescence intensity are shown on the right as in **B** (n=25). (**F** and **G**) Kap123 is required for NLS1 import into the nucleus, but not for NLS2. Expression of *csa1-(1-49)*, *csa1-NLS1-4A*, and *csa1-NLS2-4A* was under the control of the endogenous *CSA1* promoter. Line scans of fluorescence intensity are shown on the right as in **B** (n=25). (**H** and **I**) Kap95 is required for NLS2 import into the nucleus. Expression of *CSA1-GFP* and *csa1-NLS1-4A -GFP* was under the control of the *DDI2* promoter. Cyanamide was added one hour after rapamycin to induce the production of Csa1-GFP and Csa1-NLS1-4A-GFP. Line scans of fluorescence intensity are shown on the right as in **B** (n=25). At least three biological replicates were performed in the experiments presented in **B**-**I**. Scale bars, 2μm.

To pinpoint the location of additional nuclear localization sequences, we created a GFP-fused fragment containing amino acids 31 to 90 of Csa1 (Csa1-(31-90)), which localized to the nucleus and associates with the nuclear periphery (Fig. 2B). This result suggests the presence of a second NLS, which we termed NLS2, spanning the region from 31 to 90. To further investigate this possibility, we generated a GFP-fused fragment 91-221 of Csa1 (Fig. 2B). As shown in Fig. 2B, Csa1-(91-221)-GFP localized to the cytoplasm, and notably, it was distributed throughout the cytoplasm (Fig. 2B). Therefore, Csa1 appears to harbor a bipartite nuclear localization signal at its N-terminus, comprising NLS1 within 1-30 and NLS2 within 31-90.

To further demonstrate that Csa1 possessed a bipartite nuclear localization signal, we generated point mutations to NLS2 (NLS2-4A), replacing Lys39, Lys43, Lys44, and Arg45 with alanine (Fig. 2D). As with Csa1-NLS1-4A, the localization of Csa1-NLS2-4A appeared normal (Fig. 2D). However, the double mutant Csa1-NLS1-4A-NLS2-4A failed to be sorted to the nucleus, instead accumulating at cytoplasmic membrane structures (Fig. 2D). These findings collectively support the notion that Csa1 harbors a bipartite NLS signal located at its N terminus. Moreover, our results suggest that the amino acid sequence within 31-90 residues is crucial for Csa1’s association with the membrane (see below Fig. 3).

**Figure 3.**
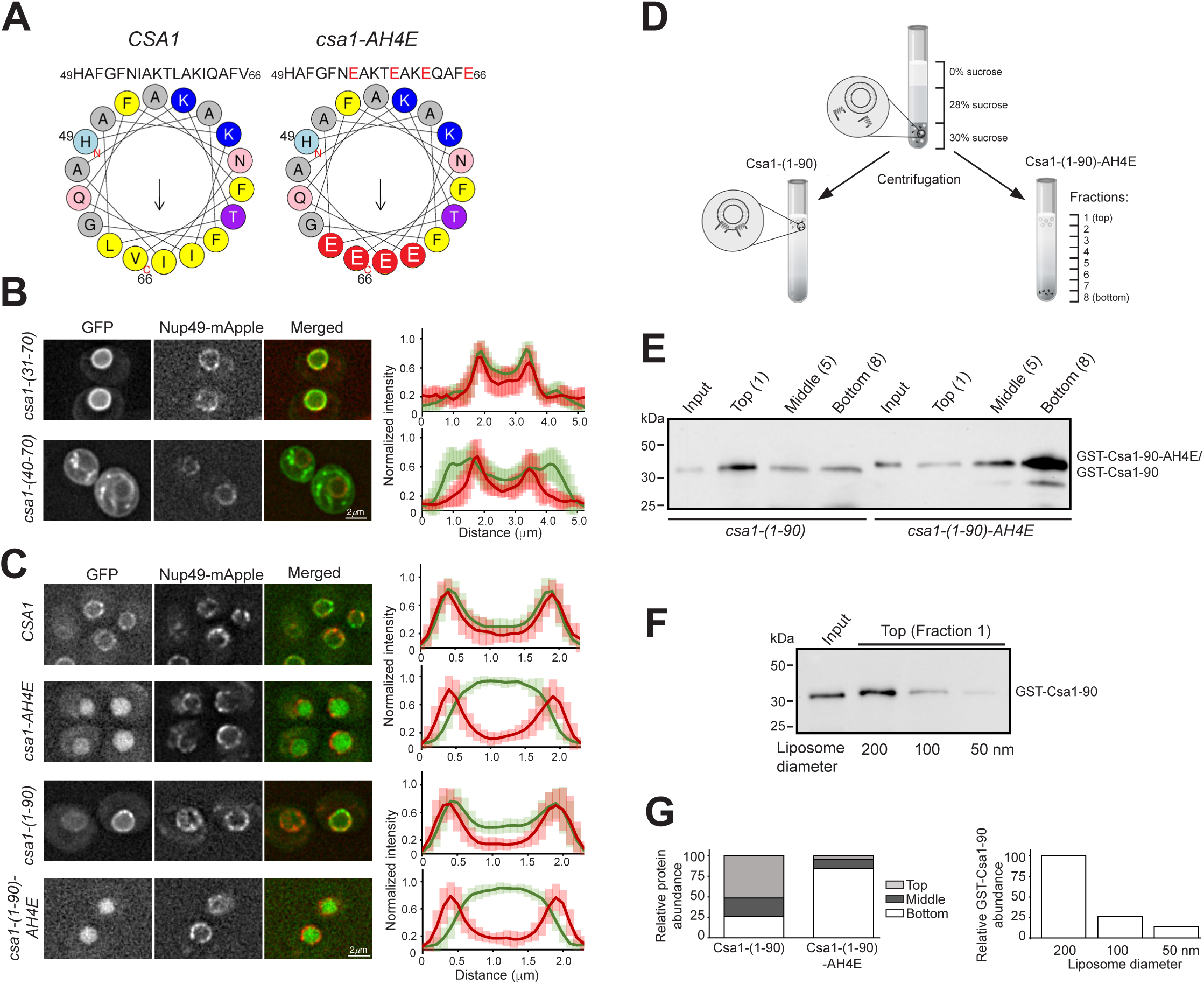
Csa1 harbors an amphipathic helix (AH) that binds to phospholipid bilayers. (**A**) Helical wheel models of Csa1-(49-66) and Csa1-(49-66)-AH4E. The positions of point mutations are shown in red. (**B**) Csa1-(40-70) is sufficient for membrane binding. Note that Csa1-(30-70) retains NLS2 and the AH domain and localizes to the nuclear envelope, Csa1-(40-70) binds to cytoplasmic membrane structures. Expression of *csa1-(30-70)-GFP* and *csa1-(40-70)-GFP* was under the control of the *DDI2* promoter. Fluorescence intensities along a 5 μm line across the cell are shown to the right (n=25). Error bars represent standard deviation. (**C**) Csa1-AH4E lacks membrane-binding activity. Expression of the full length *CSA1*, *csa1-AH4E, csa1-(1-90)* and *csa1-(1-90)-AH4E* was under the control of the endogenous *CSA1* promoter. Note that both Csa1-AH4E and Csa1-(1-90)-AH4E are localized within the nucleus. Nup49-mApple marks the nuclear periphery. Line scans of fluorescence intensity are shown on the right as in **B** (n=25). At least three biological replicates were performed in the experiments presented in **B** and **C**. (**D**) Schematic representation of the liposome flotation assay. Recombinant GST-Csa1-(1-90) and GST-Csa1-(1-90)-AH4E proteins were incubated with liposomes and subjected to centrifugation. Eight equal fractions were collected from top to bottom. Predicted outcomes of Csa1-(1-90) (left) and Csa1-(1-90)-AH4E (right) are depicted with Csa1-(1-90) cofractionating with liposomes in the top fraction and Csa1-(1-90)-AH4E accumulating in the bottom fraction. (**E**) Western blotting showing the top (1), middle (5), and bottom (8) fractions of GST-Csa1-(1-90) and GST-Csa1(1-90)-AH4E, using liposomes prepared with a 200nm pore size membrane. (**F**) Western blotting showing GST-Csa1-(1-90) in the top fraction after flotation experiments conducted using liposomes prepared with membranes of pore sizes 200nm, 100nm, and 50nm. At least three biological replicates were performed in the experiments presented in **E** and **F**. (**G**) Quantification of the Western blots shown in **E** and **F**. Left: Ratio of protein abundance between the top and bottom fractions of GST-Csa1-(1-90) and GST-Csa1-1-90)-AH4E. Right: Relative protein abundance of GST-Csa1-(1-90) cofractionating with liposomes of different diameters: 200nm, 100nm, and 50nm.

**Figure 4.**
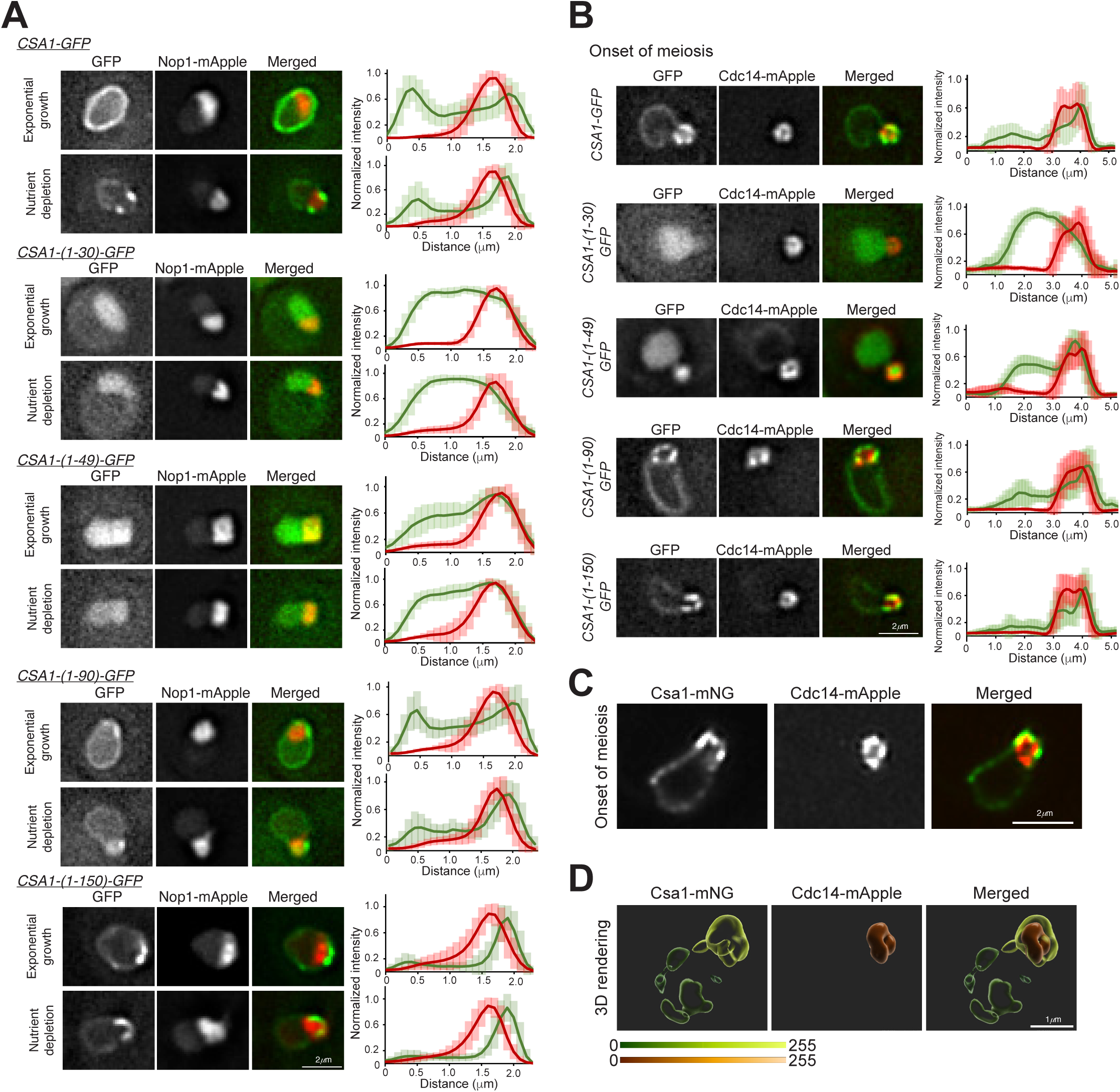
Sequence requirement for Csa1 concentration in the n-INM region. (**A**) Representative images showing the localization of Csa1 and C-terminal deletions of Csa1 during exponential growth and under nutrient depletion. The expression of all C-terminal deletions was under the control of the endogenous *CSA1* promoter. Nop1-mApple marks the nucleolus. Fluorescence intensities along a 2.5 μm line across the nucleus are shown to the right (n=25). Error bars represent standard deviation. (**B**) Representative images showing Csa1 and C-terminal deletions of Csa1 prior to meiotic entry. Fluorescence intensities along a 5 μm line across the cell are shown to the right (n=25). At least three biological replicates were performed in the experiments presented in **A** and **B**. (**C**) A single confocal section of a meiotic cell showing Csa1-mNeongreen localization by structured illumination microscopy. (**D**) Three-dimensional rendering of the cell shown in panel **C**. Heatmap depicts average fluorescence intensity of each surface. Note that Cdc14-mApple forms a torus shape.

### Kap123 and Kap95 are required for sorting Csa1 to the nucleus

To elucidate the mechanism of Csa1 import into the nucleus, we used two complementary approaches: mutational analysis of nonessential karyopherin genes and the anchor-away method to force karyopherin localization to the cytoplasm (Figs. 3E-3I). To narrow down the karyopherins responsible for Csa1 targeting to the nucleus, we took advantage of our affinity purification and mass spectrometry assay, which identified Kap123 and Kap95 as two potential Csa1 interacting karyopherins (Our unpublished data). Interestingly, Csa1-(1-30)-GFP lost its nuclear localization in *kap123Δ* cells (Fig. 2E). However, mutation of NLS1 (Csa1-NLS1-4A) was insufficient to alter Csa1 nuclear localization, as Csa1-NLS1-GFP was sorted to the nucleus and associated with the nuclear envelope in *kap123Δ* cells (Fig. 2E). Furthermore, Csa1-(1-49)-GFP was imported into the yeast nucleus in *kap123Δ* cells (Fig. 2F). These findings suggest that Kap123 is responsible for sorting Csa1-NLS1, but not Csa1-NLS2. Supporting this idea, Csa1-NLS2-4A-GFP was predominantly localized in the cytoplasm when Kap123 was absent (Fig. 2G). Since *KAP95* is an essential gene, we used the anchor-away approach. Upon addition of rapamycin, Kap95-FRB is sequestered to the plasma membrane through the formation of a ternary complex with the integral plasma membrane protein Pma1-2xFKBP12 (Hwang et al., 2022). Importantly, the full-length Csa1-GFP maintained its localization to the inner nuclear membrane upon the addition of rapamycin, while Csa1-NLS1-4A-GFP was found at the cytoplasmic membrane structures when nuclear Kap95 was depleted (Figs. 2H and 2I). These findings collectively support our hypothesis that Csa1 possesses two distinct nuclear localization sequences, NLS1 and NLS2, which interact with Kap123 and Kap95, respectively.

### Csa1 harbors an amphipathic helix at amino acid positions 49-66

Our deletional analysis of *CSA1* also revealed that amino acids 31 to 90 are crucial for the association of Csa1 with the INM (Fig. 2). We further narrowed down the amino acids to 31 to 70, which were sufficient for binding to the INM (Fig. 3B). To understand how Csa1 interacts with the INM, we employed a bioinformatics approach. Helical wheel analysis indicated that Csa1 possesses a potential amphipathic helix within amino acids 49 to 66 (Fig. 3A). Consequently, we constructed a Csa1-(40-70)-GFP fusion (Fig. 3B) and observed that amino acids 40 to 70 were sufficient to localize to membrane structures within the cell, including the cortical endoplasmic reticulum (Fig. 3B). Note that the CSA1-(40-70)-GFP fusion lacks a complete NLS2. To determine whether the amino acids 49 to 66 are essential for Csa1 binding to membranes, we mutated the nonpolar amino acids Ile55, Leu59, Ile62, and Val66 to the negatively charged Glu, which we designated Csa1-AH4E (Figs. 3A and 3C). Remarkably, the 4E mutation in the amphipathic helix abolished membrane association, leading to its localization to the nucleoplasm (Fig. 3C). Similarly, Csa1-(1-90)-AH4E was concentrated in the nucleoplasm as shown in Fig. 3C. Note that Csa1-(31-70) contains both the NLS2 and the AH domain, demonstrating the minimal requirement to associate with the INM (Fig. 3B). Together, these findings support the hypothesis that Csa1 contains a membrane-binding amphipathic helix at amino acid positions 49 to 66.

To confirm the amphipathic helix of Csa1’s ability to bind to phospholipids, we conducted an in vitro liposome co-flotation assay (Figs. 3D-3G). Among the Csa1 fragments that were expressed in *E. coli*, we found that GST-Csa1-(1-90) yielded soluble proteins that are suitable for the liposome flotation assay (data not shown). Using recombinant GST-Csa1-(1-90) and GST-Csa1-(1-90)-AH4E, we observed that GST-Csa1-(1-90) cofractionated efficiently with liposomes prepared with a 200nm polycarbonate filter, whereas GST-Csa1-(1-90)-AH4E did not (Figs. 3E and 3G). Furthermore, GST-Csa1-(1-90) showed a preference for large liposomes, as evidenced by its higher cofractionation rate with 200nm liposomes compared to 50nm liposomes (Figs. 3F and 3G). Taken together, these results show that Csa1 has an amphipathic helix at amino acid positions 49 to 66, which is both necessary and sufficient for Csa1 to directly bind to phospholipid bilayers.

### C-terminal deletions force constitutive association of Csa1 with the n-INM

We have shown that Csa1 forms a concentrated patch at the n-INM in response to starvation and at meiotic entry (Fig. 1). To further elucidate how Csa1 binds to the n-INM, we used the *csa1* deletional mutants described above (Figs. 2 and 3). We found that Csa1-30 localized to the nucleoplasm but was mostly excluded from the nucleolus region (Fig. 4A). In contrast, Csa1-49 localized to the nucleus but was more concentrated in the nucleolus region in nutrient-depleted cells (Fig. 4A). These phenotypes observed in vegetative yeast cells were also seen in cells at the beginning of meiosis, when the nucleolus-associated nuclear envelope formed a globular protrusion from the rest of the nucleus (Figs. 1E and 4B). We therefore conclude that the N-terminus of Csa1 is crucial for its localization to the n-INM.

To determine how Csa1’s localization to the n-INM changes during the cell cycle, we used time-lapse fluorescence microscopy of yeast cells marked with Csa1-GFP and Nop1-mApple (Figs. S2A and S2B). Before mitosis, Csa1-GFP was evenly distributed at the nuclear periphery (Fig. S2). During mitosis, Csa1 formed a focus at the poles, corresponding to the yeast spindle pole bodies (Fig. S2A, (Ord et al., 2020). However, Csa1’s concentration to the n-INM occurred only briefly during the G1 phase (Fig. S2). These findings were recapitulated in meiotic cells, where Csa1 was predominantly associated with the n-INM during meiotic interphase (Fig. 1F). Therefore, Csa1’s association with the n-INM is regulated during the cell cycle.

Previous research has demonstrated that Csa1 undergoes phosphorylation primarily at its C-terminus (Ord et al., 2020). Consequently, we generated C-terminal deletional alleles, *csa1-(1-90)* and *csa1-(1-150)*, which lack most of the known phosphorylation sites in its encoded protein. Surprisingly, both Csa1-(1-90) and Csa1-(1-150) were constitutively localized to the n-INM region in both cycling and nutrient-depleted cells, and throughout the cell cycle (Figs. 4A, 4B, and Fig. S2). In addition, the fragment Csa1-(31-90) uniformly localized to the nuclear periphery, while the fragment Csa1-(31-221) formed puncta along the nuclear periphery (Fig. 2B). Together, these results suggest that the C-terminus plays a regulatory role in Csa1’s localization to the n-INM.

To further elucidate Csa1’s interaction with the n-INM and the nucleolus, we used super-resolution microscopy and generated three-dimensional reconstructions of Csa1 and the nucleolus (Figs. 4C, 4D, and Fig. S3). The locally concentrated Csa1-mNeongreen formed a dome atop the Cdc14-mApple signal, which marked the nucleolus and appeared to be a torus (Fig. 4D and Fig. S3). Csa1 encircled the INM proximal side of the nucleolus, while it was absent from the distal side of the nucleolus, which lies within the nucleoplasm (Figs. 4C and 4D). These findings further support the notion that Csa1 is a membrane protein and that the n-INM constitutes a specialized region of the INM.

### Csa1 restricts Lro1 localization to the n-INM

The preferential localization of Csa1 to the n-INM region in starved yeast cells is similar to that of Lro1 (Fig. 5 and (Barbosa et al., 2019). Using fluorescence microscopy, we found that both Csa1 and Lro1 were concentrated to the n-INM region in nutrient depleted yeast cells and at the onset of meiosis (Figs. 5A and 5B). To reduce the degradation of Lro1 protein, we used the *asi3Δ* allele (Barbosa et al., 2019), and we observed the localization of Csa1 and Lro1 to the same n-INM region (Fig. 5B). We also observed that the n-INM region and the nucleolus underwent constant morphological changes during early meiosis, as revealed by time-lapse microscopy (Fig. 1F and Fig. S2). In approximately 50% of yeast cells blocked at early meiosis by *P_DDI2_-IME1*, we observed that Csa1 formed two symmetrical patches at the proximal contact sites between the nucleolus and the INM (Fig. 4D and Fig. S3). These observations led us to hypothesize that Csa1 constricts Lro1 localization to the n-INM region. In the absence of Csa1, Lro1 was no longer concentrated to the n-INM but instead formed multiple condensed foci along the nuclear periphery. To reliably observe the Lro1 protein localization at the onset of meiosis, we used the *P_CUP1_-LRO1-GFP* allele to overproduce Lro1 (Fig. 5D). We note that the localization of Csa1 to the n-INM remains unperturbed in the absence of Lro1 (Fig. 5C), suggesting that Csa1 acts upstream of Lro1. In addition, Lro1 maintained its normal protein levels and stability even when Csa1 was absent (Fig. 5E). Therefore, Csa1 plays a crucial role in constricting Lro1 localization to the n-INM, but not vice versa.

**Figure 5.**
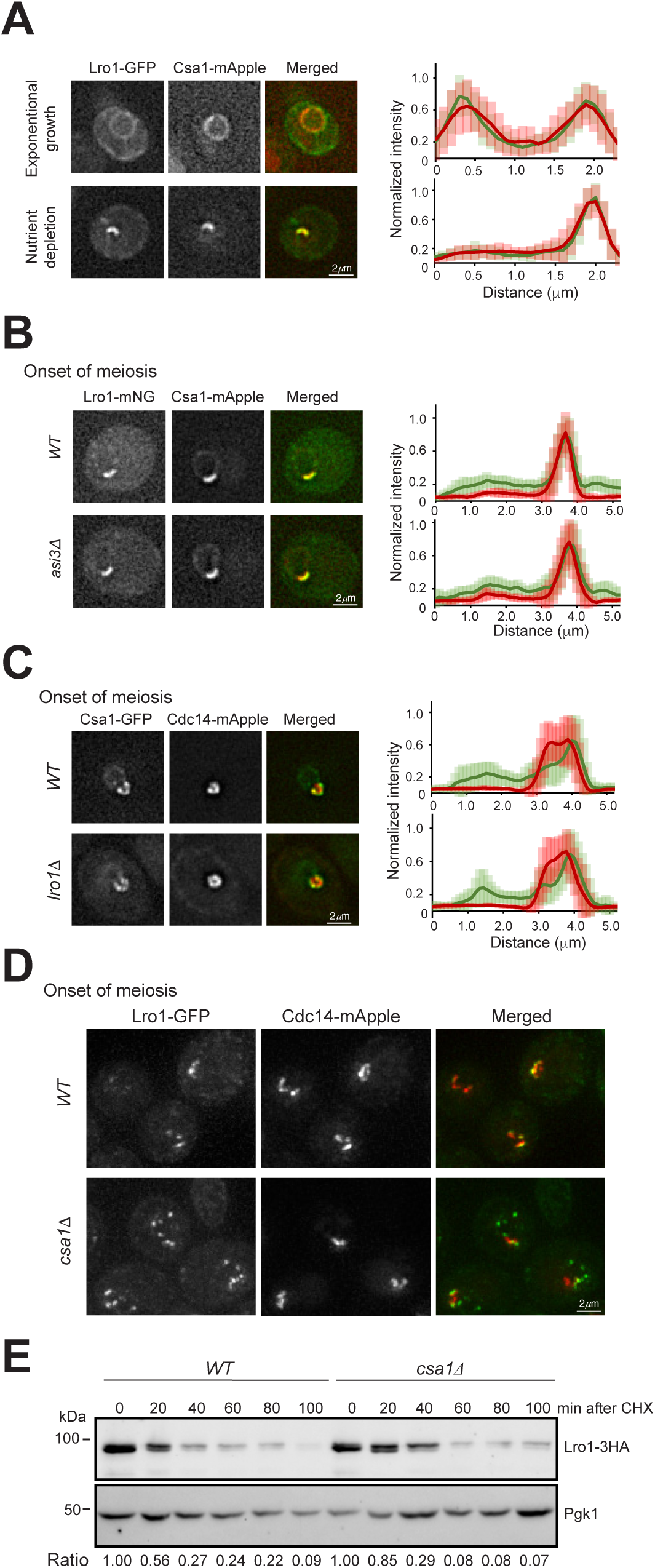
Csa1 restricts Lro1 localization to the n-INM region. (**A**) Representative images showing Csa1-mApple and Lro1-GFP localization during exponential growth and under nutrient depletion. Fluorescence intensities along a 2.5 μm line across the nucleus are shown to the right (n=25). Error bars represent standard deviation. (**B**) Representative images showing Csa1-mApple and Lro1-mNeongreen colocalization at the n-INM at the onset of meiosis. Fluorescence intensities along a 5 μm line across the cell are shown to the right (n=25). (**C**) Representative images showing that Lro1 is dispensable for Csa1 accumulation at the n-INM. Line-scans of fluorescence intensity were performed as in **B**. (**D**) Representative images showing that Csa1 is required for constricting Lro1 to the n-INM. The expression of *LRO1-GFP* is under the control of the *CUP1* promoter. Note that Lro1 forms foci at the nuclear periphery and outside the n-INM region. The displayed images are the maximum projections. (**E**) Lro1 protein stability in wild-type and *csa1Δ* cells. Yeast cells were grown to the exponential phase, then cycloheximide was added to the culture media. Cell aliquots were withdrawn at specific time points, and protein extracts were prepared for Western blotting. An anti-HA antibody was used to detect Lro1-3HA, the level of Pgk1 served as a loading control. The bottom shows the relative ratio of Lro1-3HA to Pgk1 protein abundance at indicated time. At least three biological replicates were performed in the experiments presented in **A**-**E**.

### The deacetylase Sir2 regulates the nuclear distribution of Csa1 and Lro1

To identify factors that regulate the localization of Csa1 and Lro1 to the n-INM, we used a candidate approach (Fig. 6 and Fig. S4). One of the factors we identified is the NAD+ dependent deacetylase Sir2 (Fig. 6 and Fig. S4). In starved yeast cells and cells undergoing meiosis, Csa1 was concentrated to the n-INM, while Sir2 was predominantly localized to the nucleolus, with small foci scattered around the nuclear periphery that corresponded to telomeres (Figs. 6A and 6C). Interestingly, in the absence of Sir2 or its enzymatic activity, Csa1 no longer formed a concentrated patch at the n-INM in nutrient-depleted cells. Instead, Csa1, as well as Csa1-150, localized evenly along the nuclear periphery (Fig. 6B and 6D). In addition to Sir2, we also found that Lrs4 had a minor role in Csa1 localization to the n-INM (Figs. S4A and S4B). However, in the absence of Csm1, a binding partner of Lrs4 inside the nucleolus, Csa1 localization was not perturbed (Fig. S4A). Importantly, we observed that overproduction of Sir2 suppressed the Csa1 mislocalization phenotype in *lrs4Δ* cells (Fig. S4B). Furthermore, deletion of *SIR3* or *SIR4* had no effect on Csa1 localization to the n-INM (Fig. S4C). To determine whether Cdc14 and Net1, two Sir2 interacting partners inside the nucleolus, play a role in Csa1 association with the n-INM, we performed time-lapse microscopy (Figs. S2C and S2D). We took advantage of the meiotic anaphase I, as both Cdc14 and Net1 are dissociated from the nucleolus (Li et al., 2011). Using both the *csa1-(1-90)* and *csa1-(1-150)* alleles, we observed that both Csa1-(1-90)-GFP and Csa1-(1-150)-GFP remained associated with the n-INM at anaphase I, indicating that neither Cdc14 nor Net1 is necessary for Csa1 binding to the n-INM (Figs. S2C and S2D). In addition, the regulators of rDNA tethering and silencing, including Heh1, Nur1, Fob1, and Tof2 (Kobayashi and Horiuchi, 1996; Mekhail et al., 2008), were not necessary for Csa1 accumulation at the n-INM (Fig. S4). Taken together, our findings suggest that Sir2, but not other components of the RENT complex, regulates Csa1 accumulation at the n-INM.

**Figure 6.**
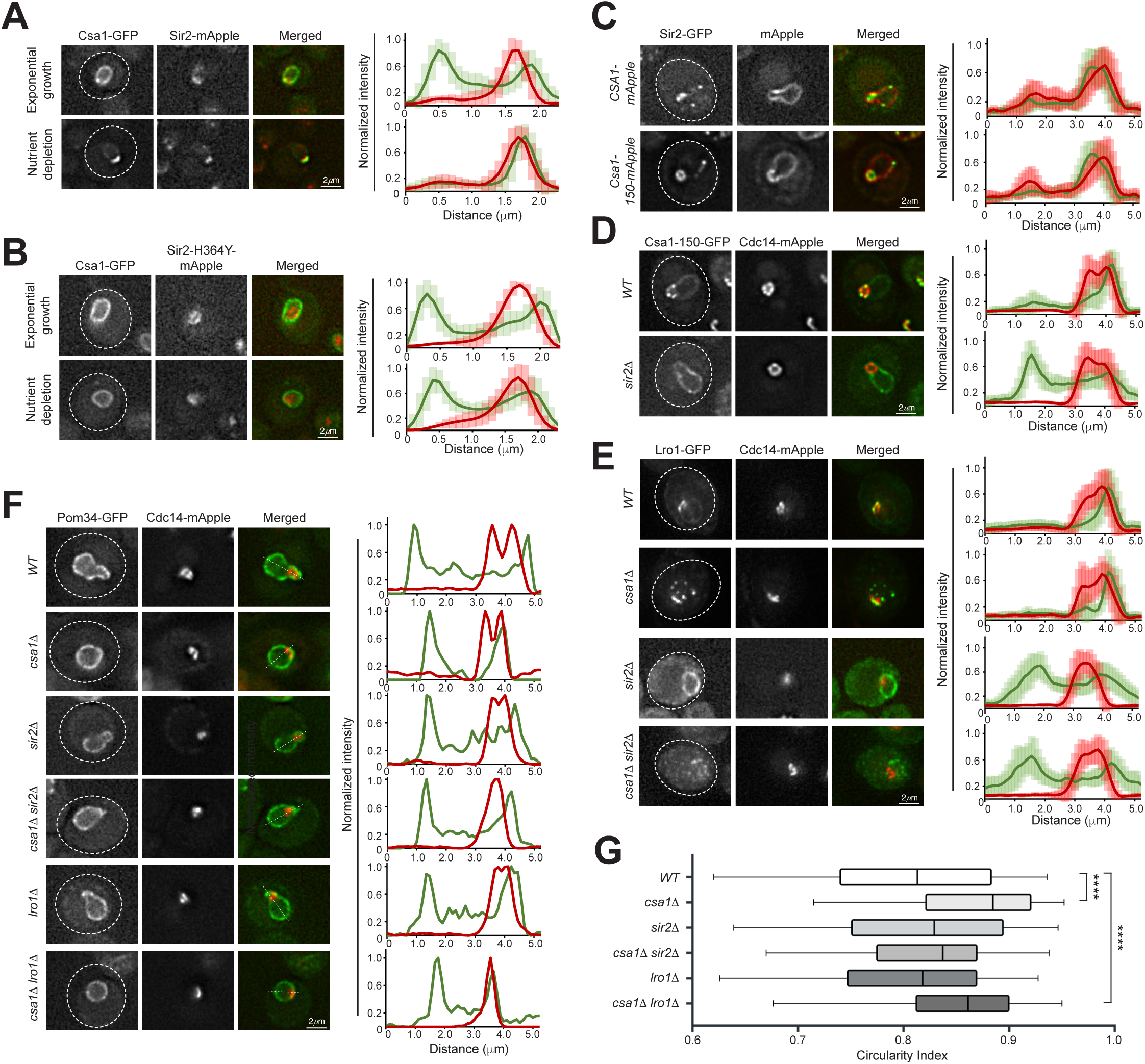
The deacetylase Sir2 regulates the nuclear distribution of Csa1 and Lro1. (**A**) Representative images showing the localization of Csa1-GFP and Sir2-mApple during exponential growth and upon nutrient depletion. Fluorescence intensities along a 2.5 μm line across the nucleus are presented on the right (n=25). Error bars represent standard deviation. (**B**) The deacetylase activity of Sir2 is required for maintaining Csa1 concentration at the n-INM. Note that *sir2-H364Y* abolishes Sir2’s deacetylase activity. Line scans of fluorescence intensity are shown on the right as in **A**. (**C**) Representative images show the localization of Sir2-GFP and Csa1-mApple and Csa1-150-mApple prior to meiotic entry. Fluorescence intensities along a 5 μm line across the cell are shown to the right (n=25). (**D**) Sir2 is required for maintaining Csa1-150 concentration at the n-INM during meiosis. Line scans of fluorescence intensity are shown on the right. In the absence of Sir2, Csa1-150 localizes evenly along the nuclear periphery. (**E**) Sir2 is required for maintaining Lro1 concentration at the n-INM. Fluorescence intensities along a 2.5 μm line across the nucleus are shown to the right (n=25), as in **A**. The displayed images are the maximum projections. (**F**) Representative images showing the nuclear morphology in *WT*, *csa1Δ*, *sir2Δ*, *lro1Δ*, *csa1Δ sir2Δ*, and *csa1Δ lro1Δ* cells. Yeast cells were arrested at the onset of meiosis using *P_DDI2_-IME1*. Pom34-GFP marks the nuclear periphery. At least three biological replicates were performed in the experiments presented in **A**-**F**. (**G**) Quantification of nuclear roundness (circularity index) in the cells shown in **F**. Bars represent standard deviation.

To determine whether the enzymatic activity of Sir2 is necessary for Csa1 localization, we used the catalytically inactive *sir2-H364Y* mutant (Tanny et al., 1999)(Fig. 6B). The enzymatically dead Sir2 protein remains localized to the nucleolus (Fig. 6B). In *sir2-H364Y* cells, Csa1 localized relatively evenly along the nuclear periphery, lacking concentration to the n-INM (Figs. 6B). This demonstrates that Sir2’s enzymatic activity is crucial for Csa1’s preferential localization to the n-INM. Similarly, we observed that Lro1 lost its focus at the n-INM in the absence of Sir2 (Fig. 6E). These results support the idea that Sir2’s enzymatic activity, but not its nucleolus localization, regulates both Csa1 and Lro1 nuclear localization.

### Csa1 regulates nuclear shape at the onset of meiosis

Our unexpected result that Csa1 modulates the nuclear localization of the acetyltransferance Lro1 led us to hypothesize that Csa1 plays a role in nuclear membrane organization. We focused on nuclear shape changes at the onset of meiosis because at this stage the nucleus undergoes a dramatic morphological change to alter cell fate (Fig. 6F). At meiotic entry, the nucleolus was extended from the rest of the nucleus (Figs. 1E and 4B), reminiscent of the mitotic nuclear flare reported previously (Witkin et al., 2012). However, we note that this meiotic nucleolus extension is phenotypically and mechanistically different from the mitotic flare. Csa1 was not concentrated at the nuclear flare, and the nuclear flare remained prominent in *csa1Δ* cells arrested in mitosis by nocodazole (Fig. S5). These observations suggest that Csa1 is not essential for mitotic nuclear flare formation. To quantify changes in nuclear shape, we measured nuclear circularity in *P_DDI2_-IME1* blocked meiotic cells. In *WT* cells, the circularity of the nucleus was 0.80, while in *csa1Δ* cells the average circularity was 0.86 (p value <0.001) (Fig. 6G). Most importantly, in *csa1Δ* cells, the nucleus lacked the nucleolus extension (Fig. 6H). Unexpectedly, *sir2*Δ could partially rescue the nuclear circularity caused by *csa1Δ,* but *lro1Δ* did not (Fig. 6F and 6G). We therefore conclude that Csa1 regulates nuclear shape prior to meiotic entry in a Sir2-dependent manner.

### Csa1 and Sir2 regulates nuclear lipid droplet biogenesis

That Sir2 modulates the nuclear localization of Csa1 and Lro1 prompted us to hypothesize that Sir2 plays a role in nuclear lipid droplet biogenesis and nuclear shape change. As Csa1 regulates the localization of the diacylglycerol acyltransferase Lro1 (Fig. 5), we sought to determine whether this change in localization results in a consequence to lipid droplet formation. Lipid droplet formation is carried out by four functionally redundant enzymes: Lro1, Dga1, Are1, and Are2 (Oelkers et al., 2002; Sandager et al., 2002). Therefore, to distinguish the relationship between Csa1 and Lro1, we measured lipid droplet formation in *dga1ΔareΔ1are2Δ* (3Δ) cells. To visualize lipid droplets, we used the neutral lipid sensor BODIPY to probe the formation and size of lipid droplets (Fig. 7A and 7B). We found that individual deletions of *CSA1* and *SIR2* resulted in a dramatic decrease in lipid droplet formation (p value < 0.001) (Fig. 7B). The combined *csa1Δsir2Δ* strains resulted in a similar decrease in lipid number, however there was no significant difference in lipid droplet formation among the *sir2Δ*, *csa1Δ*, and the combined *sir2Δcsa1Δ* mutant, suggesting that both Csa1 and Sir2 act in the same pathway. Taken together, these results support the idea that Csa1 and Sir2 regulate Lro1 localization to the INM and thereby modulates nuclear lipid droplet biogenesis.

**Figure 7.**
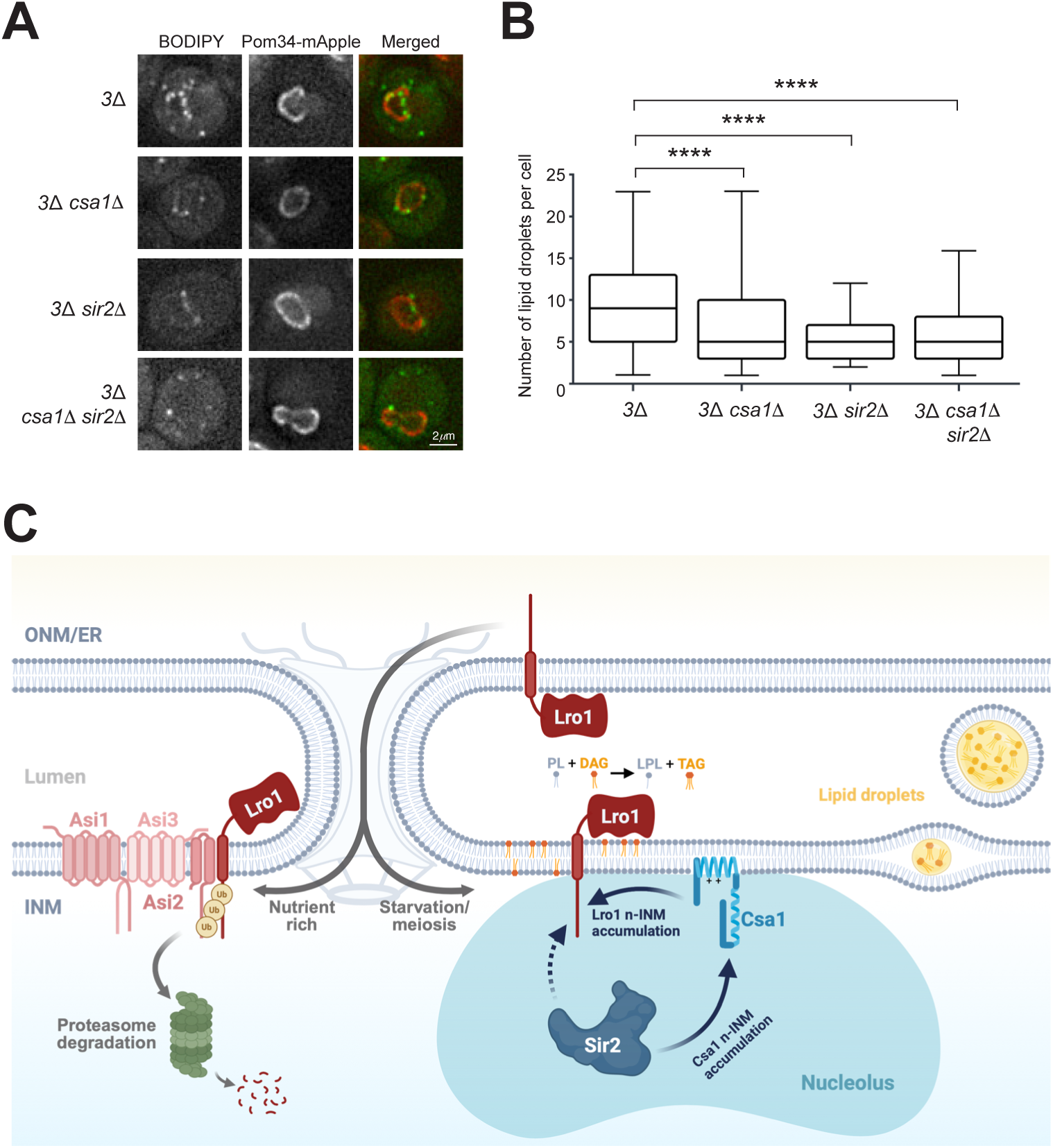
Csa1 and Sir2 regulate nuclear lipid droplet biogenesis. (**A**) Representative images show yeast cells stained by BODIPY prior to meiotic entry in *3*Δ (*are1Δ are2Δ dga1Δ*)*, 3Δcsa1Δ, 3Δsir2Δ,* and *3Δcsa1Δsir2Δ* cells. Pom34-mApple marks the nuclear periphery. (**B**) Quantification of the number of lipid droplets per cell in the strains shown in **A** (n > 200 cells per strain). (**C**) A model depicting the signaling pathway involving the Sir2-Csa1-Lro1 axis, which regulates the biogenesis of nuclear lipid droplets. In the presence of nutrients, Lro1 is imported into the nucleus and subsequently ubiquitinated by the Asi1-3 complex, leading to its degradation by the proteasome. This prevents Lro1 from accumulating at the inner nuclear membrane (INM). Conversely, when nutrients are absent or during meiotic entry, Sir2 promotes the accumulation of Csa1 at the n-INM. Upon nuclear import of Lro1, it is retained at the n-INM by Csa1, resulting in the concentration of Lro1 and the subsequent synthesis of nuclear lipid droplets. The schematic diagram was created with BioRender.com.

## Discussion

In this report, we have uncovered a signaling pathway at the INM, by which Csa1 constricts the acyltransferase Lro1 to the n-INM region in a Sir2 dependent manner. The NAD^+^-dependent deacetylase Sir2 acts as an upstream sensor that detects cellular nutrient levels. Sir2 regulates the nuclear localization of Csa1, which serves as an intermediate at the INM to recruit the effector enzyme Lro1. A resident ER membrane protein, Lro1 is imported into the nucleus upon starvation and the onset of meiosis. Csa1 constricts Lro1 to the n-INM region for lipid droplet synthesis (Fig. 7C). This reasoning is consistent with our findings that Lro1-dependent lipid droplets fail to accumulate in the absence of either Csa1 or Sir2. Our findings therefore uncover an unexpected role of Sir2 in controlling nuclear lipid droplet biogenesis, providing insights into the mechanisms of cellular aging and rejuvenation regulated by this highly conserved histone deacetylase.

Several lines of evidence we have obtained suggest that the deacetylase activity of Sir2 is required for regulating both Csa1 and Lro1 localization to the n-INM. First, in the presence of Sir2-H364Y, which abolishes the deacetylase activity, but maintains its nucleolus localization (Hoppe et al., 2002), Csa1 and Lro1 are no longer enriched at the n-INM. Instead, they are evenly dispersed along the nuclear periphery in *sir2-H364Y* cells. Therefore, localization of Sir2 to the nucleolus is not sufficient for Csa1 and Lro1 enrichment at the n-INM, but its deacetylase activity is. Second, C-terminal deletions of Csa1 constitutively bind to the n-INM region during the cell cycle and remains n-INM bound during meiotic anaphase I, when both Cdc14 and Net1 are released from the nucleolus. These findings demonstrate that Cdc14 and Net1, components of the RENT complex that recruit Sir2 to the nucleolus (Straight et al., 1999; Huang and Moazed, 2003), are not required for Csa1 binding to the n-INM. Third, Csa1 maintains its association with the n-INM in the absence of known regulators of rDNA tethering and silencing Heh1, Nur1, Lrs4 and Tof2, further supporting the idea that Sir2-dependent localization of Csa1 to the n-INM is independent of rDNA silencing. Finally, Csa1 remains enriched at the n-INM in the absence of either Sir3 or Sir4, two known Sir2 binding factors that regulate silencing at the telomeres and the cryptic mating loci (Aparicio et al., 1991). Together, our findings suggest that gene silencing at either the rDNA or the telomere is not required for Csa1 association with the n-INM. Our finding therefore differs from a recent report that telomere silencing mediated by the SIR complex is regulated by nuclear lipids (Sosa Ponce et al., 2023). We speculate that Csa1 is a potential substrate of Sir2. Consistent with this idea, Csa1 is positively charged with an isoelectric point of 10.55 and is putatively acetylated (www.yeastgenome.org). Future studies will determine the mechanism by which Sir2’s deacetylase activity regulates Csa1.

Csa1 contains two N-terminal nuclear localization sequences, NLS1 and NLS2, which interact with Kap123 and Kap95, respectively, for its nuclear import. We have revealed that the N-terminus of Csa1 is also required for INM localization, whereas the C-terminus plays a regulatory function. Deletion of the C-terminus, which removes most of the known Csa1 phosphorylation sites, renders Csa1-90 as well as Csa1-150 enrichment and constitutive binding to the n-INM. Our results therefore suggest that phosphorylation and likely additional modifications at the C-terminus have a negative effect on the enrichment of Csa1 at the n-INM.

Csa1 is enriched at the n-INM, forming dense patches that encircle the nucleolus at the onset of meiosis (this study). Combining in vivo and in vitro assays, we have shown that the AH domain of Csa1 mediates its association with the phospholipid bilayers. In addition, our lipid flotation assay shows Csa1 prefers large liposomes. Precedence exists, e.g. the basket nucleoporin Nup60, of which the AH domain introduces membrane curvatures at the nuclear envelope (Meszaros et al., 2015). We show that the nucleus is highly deformed at the onset of meiosis, where the nucleolus protrudes outward, forming high membrane curvature at the distal end (the n-INM region). In the absence of Csa1, the yeast nucleus becomes globular, consistent with the idea that Csa1 alters nuclear envelope curvature. The n-INM-bound Csa1 appears to be enriched at the proximal contact site between the INM and the nucleolus, the latter of which is protruded from the rest of the nucleus upon the induction of meiosis. Our finding therefore suggests that Csa1 is at the right location to mediate INM curvature that encircles the nucleolus, potentially establishing an INM lateral diffusion barrier that separate the n-INM from the rest of the nuclear envelope and leading to the constriction of Lro1 to this n-INM region. This reasoning is also supported by our finding that the Asi1-3 protein complex, an E3 ubiquitin ligase that degrades Lro1, is excluded from the n-INM region in a Csa1-dependent manner (see discussion below and our unpublished data). The n-INM region therefore resembles a phase-separated domain within the inner nuclear membrane. We note that the n-INM region at the onset of meiosis morphologically resembles the mitotic nuclear flare, but the physiological conditions of the cell at these two stages are completely different. Loss of Csa1 leads to globular nuclear shape at the onset of meiosis, whereas Csa1 seems to be dispensable for mitotic flare formation (this study). Curvature-forming and stabilizing proteins are often found at the edges of membrane tubules and sheets (Shibata et al., 2009; Wang et al., 2021). Our findings suggest that Csa1 is a type of such protein. In the absence of Csa1, the protruding edge at the n-INM tubule transforms into a regular sheet-like membrane structure, thereby rendering a round nucleus.

Lro1 is a resident ER protein but is imported into the yeast nucleus upon starvation and at the onset of meiosis. Current view posits that orphaned ER-associated membrane proteins, such as Lro1, are erroneously sorted to the nucleus, therefore become substrates of INMAD (inner nuclear membrane-associated degradation) and are destined for degradation by the ubiquitin proteosome system (Foresti et al., 2014; Khmelinskii et al., 2014; Koch and Yu, 2019). Indeed, Lro1 has been shown as a candidate of the Asi1-3-mediated INMAD pathway (Barbosa et al., 2019). However, upon starvation and at the onset of meiosis, Lro1 accumulates at the n-INM (Barbosa et al., 2019). Accumulation of Lro1 at the n-INM is promoted by INM-localized Csa1 and requires the deacetylase activity of Sir2 (Fig. 7C). In the absence of Csa1, Lro1 loses its enrichment at the n-INM, demonstrating that Csa1 acts as a nuclear protector of Lro1. The sanctuary provided by Csa1 at the n-INM permits Lro1 to convert DAG to TAG for lipid droplet synthesis. Our findings therefore challenge a prevalent view that INMAD substrates are erroneously imported into the nucleus only to be degraded by the proteosome. We speculate that similar orphaned ER proteins, in particular these lipid metabolic enzymes such as Lro1, are deliberately sorted to the nucleus and therefore play critical roles inside the nucleus to meet the shifting needs of the cell in response to cell fate change, such as the onset of meiosis, or environmental cues, such as nutrient depletion. Our findings also indicate that Sir2 and Csa1 are putative regulators of the INMAD pathways.

Diploid yeast cell enters meiosis in response to starvation, the process of which presents a conundrum for the cell to prepare for two continuous cell divisions with little nutrient to be acquired from its environment. Stored energy in the form of nuclear lipids therefore provides a logical energy source to initiate gametogenesis. To harvest nuclear lipids, ER-associated Lro1 is imported into the nucleus, and then sequestered to the n-INM, the process of which is aided by Csa1. Locally concentrated Lro1 and likely additional factors facilitate the production of nuclear lipid droplets, which can be transported to and consumed by the mitochondria for energy production. Similarly, nuclear lipid droplets can also be transported to the ER and participate in membrane biogenesis (Lysyganicz et al., 2025). The pathways to nuclear lipid droplet formation may differ between fungi and animals (Romanauska and Kohler, 2018; Soltysik et al., 2019). However, an emerging theme suggests that its biogenesis plays a crucial role in nuclear dynamics, beyond its function as an energy storage. The work presented here therefore provides insights into the regulation of meiotic onset and maintenance of cellular homeosis in budding yeast and beyond.

## Materials and Methods

### Yeast strains, primers and plasmids used in this study

Yeast strains used in this study and their genotypes are shown in Table S1. For mitotic experiments, strains in the S288C background were used, while SK1 strains were used for meiotic studies. Gene deletion strains were obtained from the ATCC deletion collection (ATCC-GSA-5) or through PCR-based gene deletion using homologous recombination. We used a CRISPR-Cas9-based method to generate the *sir2-H364Y* allele, which has been previously reported (Hoppe et al., 2002). We used a PCR-based homologous recombination method (Longtine et al., 1998) to tag proteins of interest to their C-terminus with GFP, mApple, mNeongreen, 3HA and V5. A similar PCR-based method was used to construct *P_DDI2_-IME1*, wherein the *IME1* promoter was replaced by the *DDI2* promoter. The sequence information of the PCR primers used in this study is provided in Table S2. Alleles that encode Cdc14-mApple, Nup49-mApple, and Nop1-mApple have been previously reported (Koch et al., 2019; Koch et al., 2020). The quadruple deletion strain (Δ4: *are1Δ*, *are2Δ*, *dga1Δ*, and *lro1Δ*) has been reported previously (Choudhary et al., 2020).

Plasmids used in this study and their key features are presented in Table S3. The split GFP plasmids have been previously described (Smoyer et al., 2016), as well as the Ycplac33-Lro1-GFP plasmid (Barbosa et al., 2019). The *DDI2* promoter (∼928bp upstream of the *DDI2* open reading frame) was amplified by PCR from BY4741. All plasmids used in this study were confirmed through DNA sequencing.

### Yeast culture methods

#### Mitotic cultures

Yeast cells were grown in synthetic complete medium (Guthrie and Fink, 1991) at 30°C. To induce the expression of the *DDI2* promoter 1 mM cyanamide (final concentration) was added after one doubling time, approximately when the optical density (OD=λ600 nm) reached 0.5 (Li et al., 2015a). To induce cells to undergo the post-diauxic shift, cells were grown from an OD of 0.2 to an OD of 4.0. To inactivate the TORC1 pathway, cells were grown from an OD of 0.1 to an OD of 0.2 before adding 250ng/mL rapamycin to the culture medium and incubating for an additional three hours prior to imaging (Fig. 1D). To deplete nuclear Kap95-FRB, we followed a previously established method (Hwang et al., 2022). To induce the expression *P_DDI2_-csa1-NLS1-4A*, we added 1 mM cyanamide to the culture medium one hour after the rapamycin (1mg/mL) treatment.

#### Meiotic cultures

Yeast cells were grown in YPD (1% yeast extract, 2% peptone, and 2% dextrose) at 30°C. To induce meiosis, the YPD cultures were diluted with YPA (1% yeast extract, 2% peptone, and 2% potassium acetate) to reach an optical density (OD) of 0.2. The diluted cultures were then incubated at 30°C for approximately 14 hours until the OD reached around 1.6–1.8. Afterward, the yeast cells were washed once in water and resuspended in 2% potassium acetate to initiate meiosis. To release cells from the *IME1* arrest in strains that express *P_DDI2_-IME1*, 500μM cyanamide was added to the culture immediately after transferring the cells to 2% potassium acetate.

### BODIPY stain of neutral lipids

Lipid droplets in both mitotic and meiotic cells were stained with 1.25 μg/ml BODIPY 493/503 for 10 minutes at room temperature before microscopy.

### Fluorescence microscopy

Epifluorescence microscopy was performed on a DeltaVision imaging system (GE Healthcare Life Sciences) at 30°C. We used a 60x (NA = 1.40) objective lens on an inverted microscope (IX-71, Olympus). Microscopic images were captured using a CoolSNAP HQ2 charge-coupled device camera (Photometrics). The pixel size was set to 0.10700 μm. Optical sections were set to 12, each with a thickness of 0.5 μm. For GFP, the excitation spectrum was at 470/540 nm, and the emission spectrum was at 525/550 nm. For mApple, the excitation spectrum was at 572/635 nm, and the emission spectrum was at 632/660 nm. To minimize photo toxicity to the cells and photo bleaching of fluorophores during time-lapse microscopy, we used neutral density filters to limit the excitation light to 50% or less of the normal equipment output. As previously described, we used agarose pads on a concave slide. A small aliquot of yeast cells was placed on the agarose, sealed with a coverslip, and then placed under microscope (Li et al., 2015b).

A Nikon CSU-W1 Spinning Disk Confocal microscope was used to collect confocal and structured illumination microscopy images. We used a 100x (NA = 1.40) objective lens on an inverted Nikon Ti2-E Motorized Inverted Microscope with Perfect Focus System. Microscopic images were acquired using an iXon Life 888 EMCCD. Super resolution images were acquired using a Live-SR module (GATACA Systems). Optical sections were set to 31, each with a thickness of 0.2 μm, using a NIDAQ Piezo Z (Andor). For GFP, the excitation spectrum was at 488 nm, and the power was set to 30%. The exposure time was 100 ms. For mApple, the excitation spectrum was at 561 nm, and the power was set to 21.3%. The exposure time was also 100 ms.

### Microscopy data analysis

Acquired DeltaVision microscopy images were deconvolved using the SoftWorx package (GE Healthcare Life Sciences). Single-section images were used for figure display unless otherwise specified. Images were then analyzed using FIJI. Linescans were generated by drawing a 1-pixel-wide line 2.5 or 5 μm in length centered at the nucleus. The plot profile function was then used to generate the mean gray value along the line. If applicable, linescans were oriented with the nucleolus closest to the end of the line. Fluorescence intensity data was normalized by first subtracting the lowest value within each linescan and then dividing by the resulting highest value. The resulting linescans (n≥25) were then averaged, and the standard deviation was calculated in Excel. Graphs depicting the normalized intensity and distance in micrometers were plotted in Excel.

To quantify lipid droplet number, images were first processed using the subtract background with rolling ball method of 50 pixels before converting to 8-bit. The unsharp mask function was then used with a radius of 3 pixels and a weight of 0.7. A manual threshold of pixel intensity was then set between 100 and 255. This image was then converted to a mask using the watershed function to separate adjacent lipid droplets. To determine the number of lipid droplets per cell, an ROI was drawn around the periphery of each cell, and the analyze particles function was used to count the number of objects within each cell.

Acquired Nikon microscopy images were deconvolved in NIS elements using the Richardson-Lucy algorithm. To create 3D renderings, processed images were then converted to Imaris files. Imaris was then used to create surfaces with a surface grain size of 0.1 micrometers. Manual threshold values were set for surfaces in each channel.

### Yeast protein extraction and western blotting

For cycloheximide (CHX)-chase experiments, yeast cells were grown overnight in YPD liquid medium to saturation at 30°C. Cell cultures were diluted with YPD to reach an OD of 0.2 and incubated at 30°C until the OD reached 0.7. At this point, CHX was added to a final concentration of 200 µg/ml, which was referred to time zero. Yeast aliquots were withdrawn at specified time points for protein extraction (Koch et al., 2019). Lro1-3HA was detected using an anti-HA mouse monoclonal antibody (1:5 K dilution, brand, cat#). The level of Pgk1 was probed using a Pgk1 antibody (Thermo Fisher Scientific, cat#PA5-28,612) as a loading control. Horseradish peroxidase-conjugated secondary antibodies, goat anti-mouse (Bio-Rad, cat#1,706,516), were used to detect the proteins of interest using an enhanced chemiluminescence (ECL) kit (Bio-Rad, cat#1,705,060). The ChemiDoc MP Imaging System (Bio-Rad, cat#17,001,402) was used to detect the ECL-based western blot. To calculate the relative protein abundance at each time point, individual band intensities were measured in mean gray value using FIJI and exported to Microsoft Excel. Target protein band intensities were normalized to those of the loading control (Pgk1).

### Recombinant protein production and purification

GST-CSA1-(1-90) and GST-CSA1-(1-90)-AH4E were cloned into the BamHI and SmaI sites of the pGEX-2T expression vector (Roche). The cloned plasmids were transformed into BL21 cells, and the expression of the recombinant proteins was induced at 22°C for 3 hours. Harvested E. coli cells were lysed in 50 mM Tris (pH 7.4) and 150 mM NaCl, supplemented with a cocktail of anti-proteases (Sigma-Aldrich). The crude lysates were centrifuged at 4°C for 10 minutes at top speed. The collected supernatants were incubated for 1 hour with the glutathione SuperFlow agarose (Thermo Fisher Scientific, CAT#25236). Before elution, the agarose beads were thoroughly washed. Finally, the GST fusion proteins were eluted in the wash buffer supplemented with 10 mM glutathione.

### Liposome preparation and flotation assay

Lipids used in this study were purchased from Avanti Polar Lipids. A 10 mM stock solution of 18:1 (Δ9-Cis) PC (DOPC) (Avanti, cat#4235-95-4,) and 18:1 (Δ9-Cis) PE (DOPE) (Avanti, cat#4004-05-1,) were resuspended in chloroform and stored at -20°C. Liposomes were prepared using protocols previously published (Kupke et al., 2011; Bhaskara et al., 2019). Briefly, on the day of the experiment, 360 μg of DOPC:DOPE (80:20) was measured out and transferred into a 1.5 ml Eppendorf tube. The tube was then dried in a speed vacuum for 30 minutes at room temperature. The dried liposome suspension was then resuspended in 360 μl of 50 mM Hepes (pH 7.2) and 150 mM NaCl (liposome buffer). After dehydration, the liposome suspension was sonicated (VWR, cat#B3500A-MTH) and then rehydrated. The liposome suspension was then extruded sequentially through 400, 200, 100, and 50 nm polycarbonate filters (Cat#610007, 610006, 610005, and 610003, respectively) using a hand extruder (Avanti, cat#610000,) at a final lipid concentration of 10 mg/ml. The liposomes were used immediately after preparation.

We followed a previous protocol for the flotation assay (Kupke et al., 2011). Briefly, recombinant proteins (60 mg) and liposomes (600 μg) were incubated in liposome buffer at 37°C for 2 hours in a total volume of 120 μl. The suspension was adjusted to 30% sucrose by adding and mixing 145 μl of a 60% w/v sucrose solution in the liposome buffer. The resulting high-sucrose suspension was overlaid with 400 μl liposome buffer containing 28% w/v sucrose and 135 μl liposome buffer containing no sucrose. The sample was centrifuged at 115,000g in a TLA-110 rotor (Beckman) at 20°C for 2 hours. Eight 100 μl fractions were manually collected from the top to the bottom. Binding affinity was determined by Western blotting using a GST antibody (Invitrogen CAT#PA1982, 1:5000).

## Supplemental Tables

Table S1: Yeast strains used in this study

Table S2: Primers used in this study

Table S3: Plasmids used in this study

Table S4: Critical reagents and antibodies used in this study

**The following sites were utilized for protein alignment and helical wheel analysis.** https://www.ebi.ac.uk/jdispatcher/psa/emboss_needle https://heliquest.ipmc.cnrs.fr/index.html

## Acknowledgments

We thank S Lenhert, R Tomko, Y Wang, and the Yu lab members for their invaluable discussions. S Lenhert provided technical support in the preparation of liposomes. We also express our gratitude to the Prinz lab, the Siniossoglou lab, and the Lusk lab for providing yeast strains and plasmids. A. Kalpetta Gramathil assisted in creating the *P_DDI2_-IME1* strain. Additionally, we would like to thank the Biological Science core facility for their assistance in DNA sequencing and plasmid construction, and the Biological Imaging facility for access to the Nikon CSU-W1 Spinning Disk Confocal microscope. This research was supported by the National Institutes of Health (GM138838) and the National Science Foundation (MCB1951313).

## Supplemental figure legends

**Figure S1.**
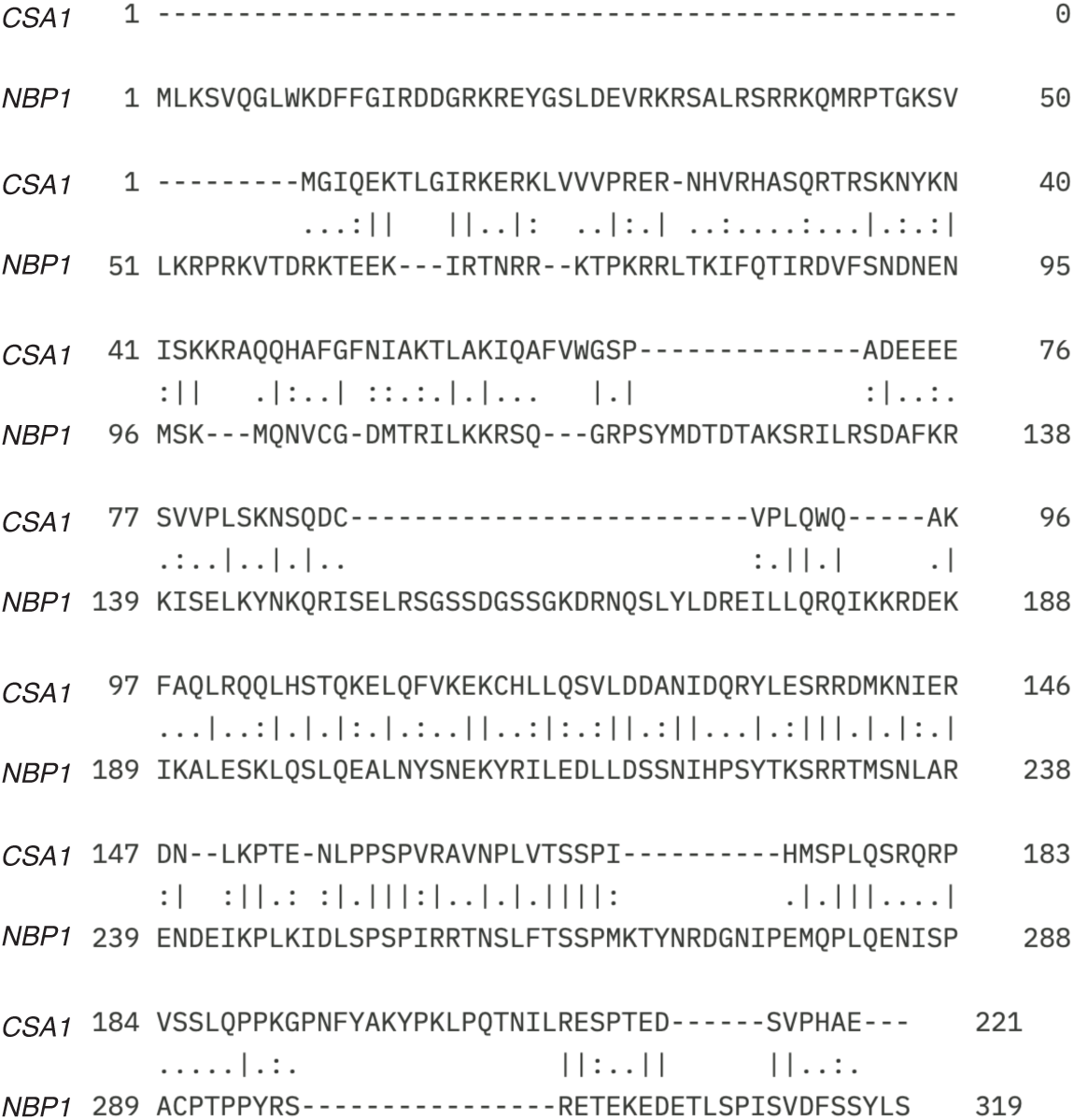
Pairwise sequence alignment of *CSA1* and *NBP1*. Identical amino acids are indicated by “|”, conserved amino acids by “:”, and semi-conserved amino acids by “.”

**Figure S2.**
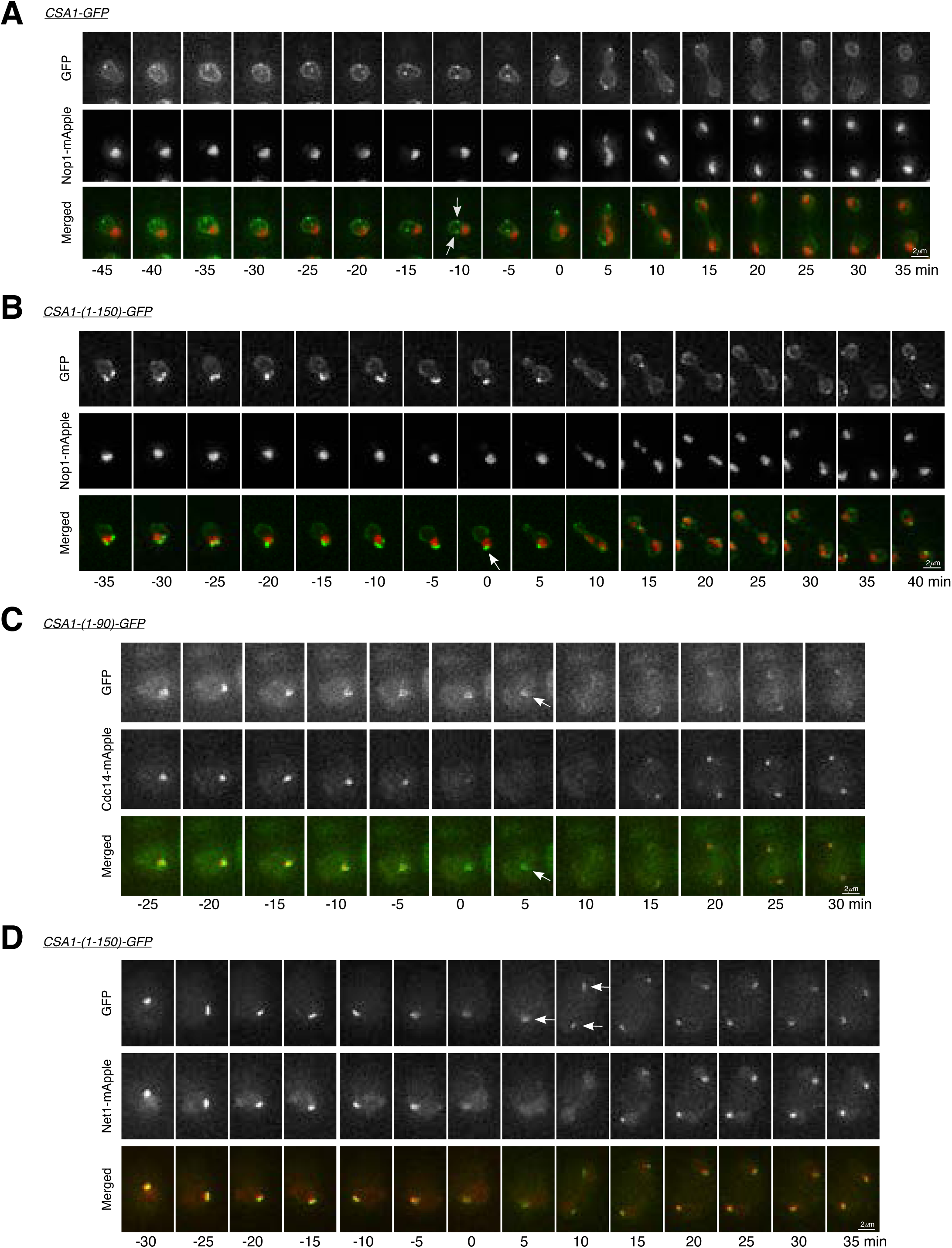
Time-lapse fluorescence microscopy shows the localization of Csa1-GFP, Csa1-(1-90)-GFP, and Csa1-(1-150)-GFP in dividing mitotic and meiotic yeast cells. (**A**) Csa1-GFP localization in a dividing mitotic cell. Arrows point to the concentration of Csa1-GFP at the spindle pole bodies. Nop1-mApple marks the nucleolus. Time zero refers to the onset of anaphase. (**B**) Localization of Csa1-(1-150)-GFP in a dividing mitotic cell as shown in **A**. The arrow points to the concentration of Csa1 to the n-INM region. Time zero is also at the onset of anaphase. (**C**) Localization of Csa1-(1-90)-GFP in meiosis I. Arrows point to the concentration of Csa1-(1-90)-GFP at the n-INM region at anaphase I. Time zero is at the onset of anaphase I. (**D**) Localization of Csa1-(1-150)-GFP in meiosis I as shown in **C**. Arrows point to the concentration of Csa1-(1-150)-GFP at the n-INM region at anaphase I. Net1-mApple marks the nucleolus. Time zero is at the onset of anaphase I. During anaphase I, both Cdc14-mApple and Net1-mApple are dispersed. The displayed images are the maximum projections.

**Figure S3.**
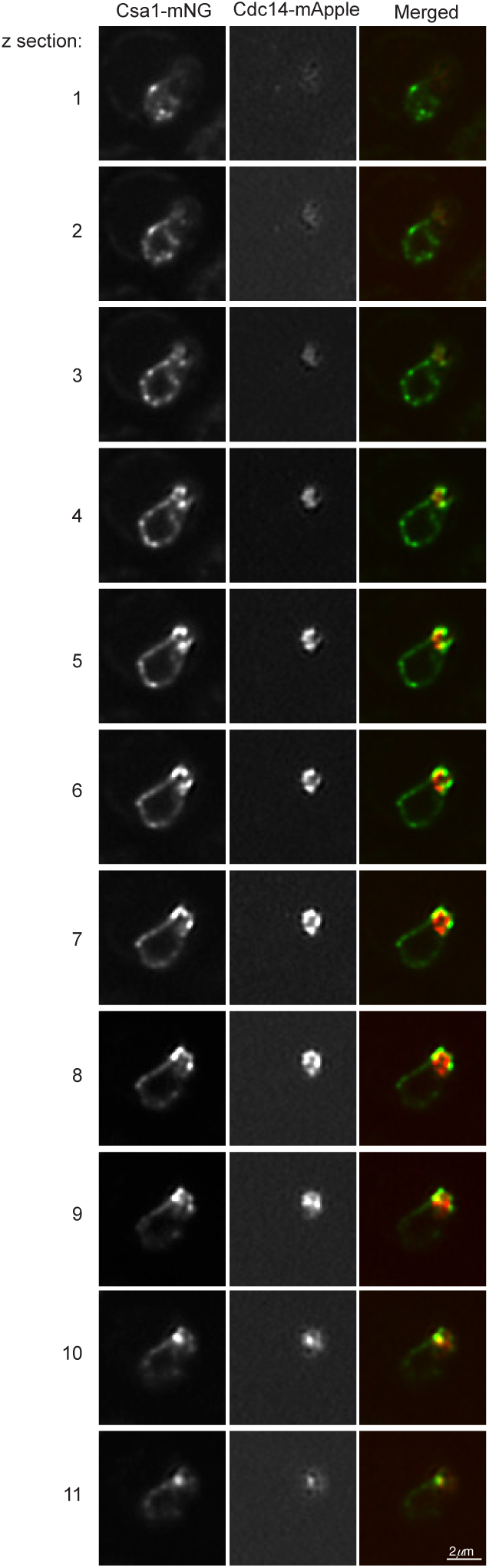
Z-series of Csa1-mNG localization prior to meiotic entry. Cdc14-mApple marks the nucleolus. Step size = 0.2µm.

**Figure S4.**
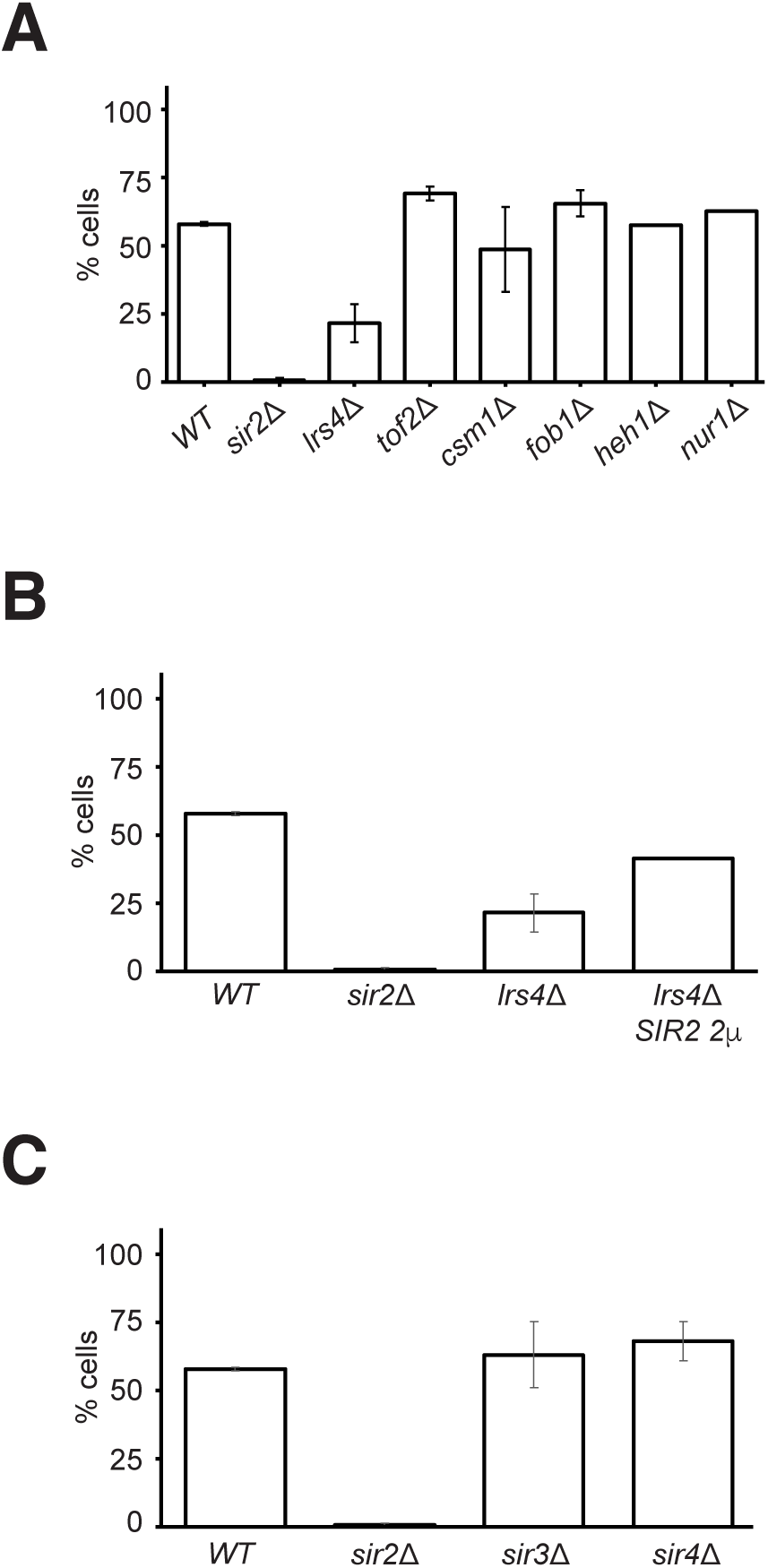
Nuclear factors regulate the localization of Csa1 to the n-INM region. (**A**) Quantification of Csa1-GFP concentration at the n-INM in starved yeast cells from the following strains: *WT*, *sir2Δ, lrs4Δ, csm1Δ, tof2Δ*, and *fob1Δ*. Data is derived from three biological replicates, each containing more than 50 cells per strain. Error bars represent standard deviation. **(B)** Quantification of Csa1-GFP concentration at the n-INM in starved cells from the following strains: *WT*, *sir2Δ, lrs4Δ* and *lrs4Δ SIR2* 2µ (n>50). Note that *SIR2* 2µ overproduces the Sir2 protein. **(C)** Quantification of Csa1-GFP concentration at the n-INM in starved yeast cells from the following strains: *WT, sir2Δ, sir3Δ,* and *sir4Δ* (n>300).

**Figure S5.**
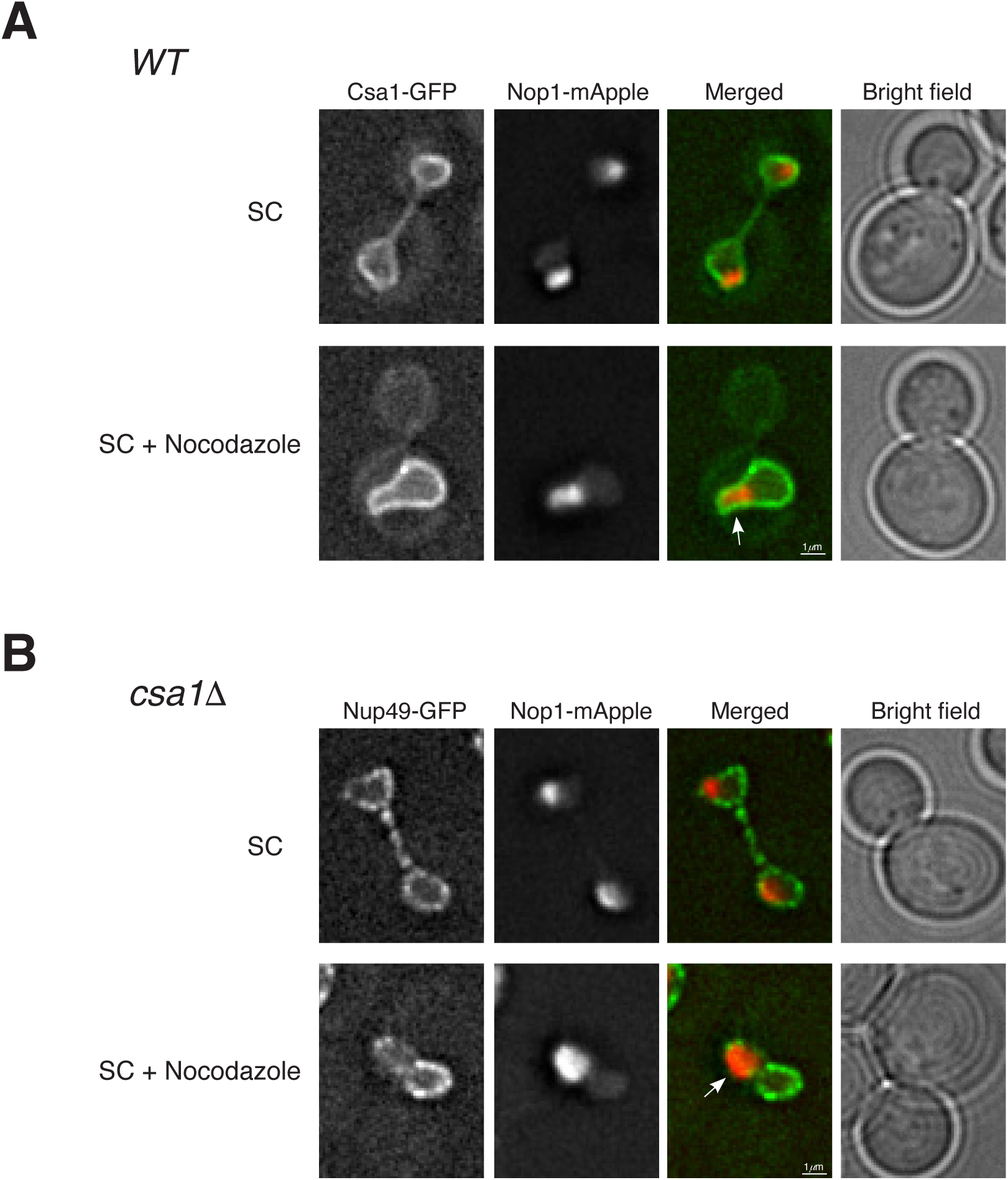
Csa1 is not required for the formation of nuclear flares in arrested mitotic cells. (**A**) Csa1-GFP localizes evenly around the nuclear periphery in a dividing (top) and an arrested (bottom) mitotic cell. Note that nocodazole arrests yeast cells at metaphase. Nop1-mApple marks the nucleolus. (**B**) The nuclear morphology of a dividing (top) and an arrested *csa1Δ* cell during mitosis. Note that nuclear flare forms in the absence of Csa1. Arrows point to the mitotic nuclear flares.

